# Coordinated leaf hydraulic thresholds maintain virtually null stomatal safety margins in poplar despite genetic variation and nutrient-induced phenotypic plasticity

**DOI:** 10.64898/2026.07.10.737750

**Authors:** D. Chassagnaud, L. Bezon, I. Le Jan, R. Fichot

## Abstract

The sequence of leaf physiological thresholds underlying plant responses to water deficit is thought to be functionally coordinated; yet, to what extent this coordination is maintained across genotypes and environments remains poorly documented at the intraspecific level. We characterized the sequence of stomatal closure, turgor loss and xylem embolism in the leaves of two genotypes of the riparian species *Populus nigra* (DRA-038 vs. PG-31) subjected to control, additional nitrogen or additional potassium treatments. Under control conditions, embolism measurements using the optical vulnerability method showed that DRA-038 was more vulnerable than PG-31, in agreement with measurements performed on stems with the reference Cavitron method. Stomatal closure consistently preceded xylem embolism, while bulk leaf turgor loss was typically observed once xylem embolism had already reached 50%. Hydraulic thresholds responded to treatments in a genotype-dependent manner, the intrinsically more vulnerable genotype DRA-038 being typically more plastic. However, despite variations across genotypes and treatments, the trait sequence remained tightly coordinated such that stomatal safety margins (SSMs) remained virtually null. These findings support a strong mechanistic integration of leaf hydraulic thresholds in poplar across genetic units and varying environments, questioning whether to favour intrinsic tolerance or plastic capacities in breeding future drought-tolerant genotypes.

## Introduction

Ongoing climate change increasingly exposes plants to water deficit across large portions of emerged lands (Xu et al. 2019). Consequences are already visible, especially for perennial woody plants with extensive forest diebacks now well documented across the globe (Anderegg et al. 2016). Such deterioration of forest health directly alters the functioning of Earth’s terrestrial ecosystems, with implications for biodiversity and ecosystem services. Identifying the suites of traits and physiological thresholds involved in drought tolerance and survival has therefore been a central focus of tree physiology over the past few decades (McDowell et al. 2008, 2022; Bartlett et al. 2016; Mantova et al. 2022).

Drought tolerance is a complex composite attribute determined by numerous underlying traits that may interact and that are involved in sequence along the progression of water deficit. In particular, the water potentials associated with core processes such as stomatal closure, the maintenance of cell turgor or the maintenance of xylem hydraulic continuity are expected to set key functional limits and impact overall plant performance under water deficit (Bartlett et al. 2012, 2016; Martin-StPaul et al. 2017). The exact timing between these processes has long been a matter of debate, especially between leaf stomatal closure and the inception of xylem embolism (see Cochard & Delzon 2013), but the development of non-destructive phenotyping approaches, as well as large-scale interspecific comparisons, have helped clarify the spatiotemporal sequence of physiological responses to dehydration (Brodribb et al. 2016a,b; Bartlett et al. 2016; Trueba et al. 2019; Jin et al. 2023; Mantova et al. 2023). At the leaf level, there is now ample evidence that stomata generally close before any significant embolism occurs although stomatal safety margins (i.e. the delay between both processes) can be variable (Brodribb et al. 2016a; Höchberg et al. 2017; Martin-StPaul et al. 2017; Blackman et al. 2019; Creek et al. 2020; Petek-Petrik et al. 2023). The positioning of leaf turgor loss regarding other processes seems to be more variable depending on species ecology and drought tolerance strategies (Brodribb & Holbrook 2003; Trueba et al. 2019; Mantova et al. 2023; Petek-Petrik et al. 2023; Ziegler et al. 2023, 2024; Schönbeck et al. 2025). Therefore, although the response sequence to dehydration is overall well described and the functional coordination between traits is supposed to prevent severe tissue damage from uncontrolled water loss, variation between species does occur in absolute trait values but also in the spacing between thresholds thus reflecting different adaptive strategies. Variation within species is also to be expected but this aspect remains historically much less documented, especially for whole traits sequences (Dayer et al. 2020, 2022). Nonetheless, addressing within-species variation can help understanding species microevolution and can have practical implications in terms of management of genetic resources and breeding.

Genetic variation can manifest phenotypically through inherent trait values but also through the extent of trait plasticity. Phenotypic plasticity, i.e. the ability of an individual genotype to express different phenotypes depending on the prevailing environment can be a key component of flexibility for sessile organisms, especially for perennials such as trees (Nicotra et al. 2010). Like any other traits, the underlying morphological and physiological drivers of tolerance to water deficit may be responsive to environmental factors such as, for instance, nutrient availability. Besides water, nutrients are major limiting resources for plant and forest productivity (Fisher et al. 2012). Nitrogen (N) has historically received much attention because of its quantitative importance and its direct benefits on growth-related processes under non limiting water. Higher N availability, if not excessive, generally stimulates photosynthesis and carbon allocation to the aerial compartment at the expense of NSC storage (Li et al. 2020a; Liang et al. 2020). Nitrogen availability also has the potential to affect a wide array of other physiological parameters related to water relations including stomatal functioning (Graciano et al. 2005; Villar-Salvador et al. 2013; Fan et al. 2022), turgor-related parameters (Graciano et al. 2005; Bucci et al. 2006; Domec et al. 2009; Fang et al. 2018; Zhang et al. 2021) and xylem hydraulics (Harvey & van den Driessche 1999; Bucci et al. 2006; Domec et al. 2009; Hacke et al. 2010; Plavcová & Hacke 2012; Zhang et al. 2021; Fan et al. 2022; Bouyer et al. 2023). In addition to N, potassium (K) is another important macronutrient involved in both carbon and water relations. Potassium is one of the most abundant cations in the cytosol and participates in pH maintenance, enzyme activation and cation/anion balance (Ragel et al. 2019). It thus has been found to influence leaf osmotic adjustment (Battie-Laclau et al. 2014a), leaf gas exchange rates or stomatal dynamics (Battie-Laclau et al. 2014a,b, 2016; Mateus et al. 2022) but also broader physiological aspects such as photosynthate loading and carbon allocation (Epron et al. 2011; Battie-Laclau et al. 2016, Mateus et al. 2022), wood formation (Ache et al. 2010) and xylem function (Harvey & van den Driessche 1999; Mateus et al. 2024). Thus, by affecting multiple aspects of plant water and carbon economy, the availability of nutrients such as N or K has the potential to condition and alter individual drought response trajectories (Gessler et al. 2017). However, depending on species or genotypes, the effects might be variable, be it in direction and/or magnitude.

If the sequence of stomatal, hydraulic and wilting responses to water deficit is overall well established and if the effects of nutrients on plant physiology are extensively documented, it is still largely unknown to what extent the functionally coordinated traits sequence involved in plant dehydration can be uncoupled at the intraspecific level by both genetic variation and phenotypic plasticity and be related to different drought response strategies. Such knowledge may be particularly valuable, especially for species of agronomic interest in a context of breeding for more drought-tolerant plant material (Dayer et al. 2022). Poplars are fast-growing and economically important trees widely used in forestry and bioenergy plantations under temperate latitudes and constitute a well-established model for the study of woody plant physiology (Jansson & Douglas 2007). As early successional species, poplars typically show considerable physiological and morphological plasticity in response to resource availability, especially water and nutrients (e.g. Street et al. 2006; Novaes et al. 2009; Plavcová & Hacke 2012). This allows them to cope with potentially rapid environmental fluctuations in their native, mostly riparian, habitat, and from a practical point of view makes them a good model to investigate environmentally induced trait variation. In Europe, the European black poplar (*Populus nigra* L.) is one of the most common and widespread poplar species. Besides ecological and patrimonial roles, *P. nigra* has an important economic value as it is one of the main parental pools used in interspecific hybrid breeding programs. Understanding the sources of phenotypic variation occurring for key functional traits in natural populations of *P. nigra* therefore has been a cornerstone in this species (Rohde et al. 2011; Guet et al. 2015; Viger et al. 2016; Fichot et al. 2024), although an integrative perspective on the key traits involved during the sequence of response to water deficit is lacking.

In this paper, we characterized the sequence of responses to water deficit at the leaf level, from stomatal closure to turgor loss and hydraulic failure, in genotypes of the riparian species *Populus nigra* originating from contrasting geographic origins and subjected to variable N or K treatments. We built upon available genetic resources in this species and we used the non-destructive optical vulnerability (OV) method to monitor embolism development. Specifically, we hypothesized (1) that trait values associated with key physiological thresholds would vary between genotypes but remain tightly integrated along the response sequence, and (2) that N and K treatments would differentially alter absolute trait values depending on genotypes (i.e. there is genetic variation in phenotypic plasticity) potentially uncoupling the response sequence. As a side objective, we also tested to what extent the OV method was able to reliably differentiate closely related genotypes in highly vulnerable species such as poplars by comparing our findings to another reference method.

## 2. Materials and methods

### 2.1. Plant material, growing conditions and experimental design

Experiments were conducted on two genotypes of European black poplar (*Populus nigra* L.). The genotype DRA-038 originated from South-East France within the Dranse river basin while the genotype PG-31 originated from Central Italy within the Paglia river basin (more details on the origin of these genotypes are available from the GnpIS Information System, Steinbach et al. 2013). These two genotypes were chosen from a larger set of six original genotypes based on growth performance under moderate water deficit and stem vulnerability to embolism (Duplan et al. 2026). Stem vulnerability curves obtained with the Cavitron method revealed that PG-31 was intrinsically more tolerant than DRA-038 [Sup. Fig. S1; Ψ_50-stem_ = −1.95 MPa (CI_0.95_ = −1.86/-2.04 MPa) vs. −1.62 MPa (CI_0.95_ = −1.52/-1.75 MPa), respectively]. The two genotypes also showed distinct physiological responses when subjected to moderate water deficit at the molecular level: the intrinsically less tolerant genotype DRA-038 exhibited a much higher plasticity in hormonal profiles and transcriptional responses than the intrinsically more tolerant genotype PG-31 (Duplan et al. 2026).

The experiment was initiated from homogeneous dormant cuttings sampled at the end of winter from current-year branches of trees growing in the same common garden (State Forest Nursery of Guéméné-Penfao, France). Cuttings 20 cm-long were rooted in 4L pots with a standard potting mix based on sod peat, wood fibres and perlite (Klassmann® 5-666) and grown in a growth chamber under non-limiting irrigation and controlled conditions [photoperiod 16/8 (day/night, h), air temperature cycles 20/15 (day/night, °C) and constant PPFD of 950 µmol.s^-1^.m^-2^]. Bud burst occurred within two weeks following potting with no difference between genotypes. Multiple shoots originating from the cuttings were rapidly and systematically pruned to leave one single dominant shoot.

After six weeks, when the saplings were *c*. 50 cm in height, plants were randomly assigned to three treatments (*n* = 6 per treatment per genotype): control (C), additional nitrogen (N) or additional potassium (K). Plants from the C treatment were fertilized twice a week with 100 ml of a standard N-P-K 20-20-20 fertilizer (0.5 g.L^-1^ dilution following the lower range of the manufacturer’s recommendations, *i.e.* 0.61 mM NH_4_NO_3_; Plant Products Co. Ltd., Leamington, ON, Canada). Plants from the N treatment were fertilized twice a week with 100 ml of the standard solution supplemented with NH_4_NO_3_ (final NH_4_NO_3_ concentration approx. ten times higher than the standard solution, *i.e.* 6.86 mM). Plants from the K treatment were fertilized twice a week with 100 ml of the standard solution supplemented with KCl (final K concentration approx. four times higher than the standard solution, *i.e.* 8.89 mM). To avoid water deficit, plants were irrigated with ultra-pure water whenever necessary on the other days. Fertilization treatments were maintained for two months before measurements started.

For all measurements, two saplings could be processed every day in parallel such that all plants (*n* = 36) were processed within less than four weeks; plants from the two genotypes and the three treatments were systematically picked up randomly to minimize potential genotype × ontogenic effects. The saplings were brought to the laboratory the evening preceding measurements and were bagged overnight with thick black plastic bags to minimize transpiration and favour the equilibrium at high water potentials before the start of measurements the morning after.

### 2.2. Pressure-volume (p-v) curves

Leaf p-v curves were generated on one healthy mature leaf per plant (*n* = 6 per genotype per treatment) sampled early in the morning from the well-watered plants equilibrated in the dark overnight. We used the bench drying technique (Sack et al. 2010). Briefly, leaves were progressively dried on the laboratory bench and we measured leaf water potential (Ψ_leaf_, MPa) and leaf mass (± 1mg) at regular intervals. Leaves were systematically allowed to equilibrate in opaque zip plastic bags for 10 min before measuring water potential using a Scholander-type pressure chamber (DG MECA, Bordeaux, France). The timings of measurements were adjusted to target 0.2 – 0.3 MPa intervals until leaves exceeded −3 MPa so that 8-10 points could be used to construct p-v curves. The bulk leaf osmotic water potential at full hydration (π_o_, MPa), the bulk leaf water potential at turgor loss point (π_tlp_, MPa), the relative water content at turgor loss point (*RWC*_tlp_, %) and the modulus of elasticity (ε, MPa) were estimated for each sample using the p-v curve-fitting routine (PVAST) provided by Sack et al. (2010).

### 2.3. Leaf vulnerability to embolism (optical method)

Leaf xylem vulnerability curves were constructed using the optical vulnerability (OV) method (Brodribb et al. 2016a). The OV method is based on the principle that light interacts differently with xylem that is water-filled *vs*. air-filled such that the subtractive analysis of image time-series allows the progressive visualization and quantification of xylem embolism over time. Embolism imaging and data storage were performed using Cavicams (University of Tasmania, Newnham, Australia). The systems consist in custom-built clamps embarking a fixed LED source and a Raspberry Pi-based time-lapse high-resolution camera allowing the capture of images at a 30× magnification of a 30 mm^2^ visible area under user-adjustable configurations of exposition, duration and interval. We installed one sensor per plant on one healthy, fully expanded leaf that had been formed during the period of treatment to make sure to account for potential treatment-induced acclimation. Sensors were installed early in the morning once plants had been debagged on the centre of the leaf blade so that images included the midrib and smaller vein orders. Measured leaves were typically located on the main stem in the top third of the canopy; when apical dominance was challenged by sylleptic branching, which can be sometimes important in poplar especially in response to nutrient fertilization, the sensor was installed on a leaf located on one of the codominant branches.

Plant dehydration was artificially generated by cutting saplings at the stem base. By repeatedly recording images of the clamped leaf from the time of cutting to complete leaf desiccation, changes in light transmission through leaf veins could be used to detect embolism in xylem conduits in time. Based on preliminary tests on our plant material, image acquisition was set to every 60 sec (shutterspeed 1700, ISO 100, max. 2592×1944 resolution) until the next morning to ensure we captured complete hydraulic failure during the recording sequence; hydraulic failure was typically reached within 8-10 hours because of the high vulnerability of poplars. In parallel to image acquisition, we measured Ψ_leaf_ repeatedly 8-10 times during the drying period using a Scholander pressure chamber on individual leaves like the ones equipped with the OV sensors. Sampled leaves were systematically allowed to equilibrate for 10 min in opaque zip plastic bags before being measured.

Image stacks were processed and analysed using automated processing scripts (CaviTools; Giese et al. 2024). Briefly, the area of embolized pixels was measured in each image using a threshold value of three and a noise reduction value of 13, which provided the best trade-off between embolism event detection and noise on our material. The cumulative embolized pixel area over time was calculated as a percentage of the total cumulative pixel area obtained at the end of the drying sequence once hydraulic failure was fully apparent in the minor vein orders. We then related each individual Ψ_leaf_ measurement (MPa) to the corresponding cumulative percentage of embolized pixels (PEP, %) based on the time precisely recorded so that a vulnerability curve could be obtained for each individual plant (Mantova et al. 2023).

### 2.4. Stomatal conductance and timing of stomatal closure

To evaluate the timing of stomatal closure during drying, we continuously measured leaf stomatal conductance to water vapour (*g*_s_, mmol.m^-2^.s^-1^) on the same plants that were measured for leaf embolism. Measurements were performed using open gas exchange systems (Licor 6400 and Licor 6800, LI-COR, Lincoln, NE, USA) on a healthy, fully mature leaf located close to the OV sensor with the following parameters: reference atmospheric [CO_2_] of 400 ppm, saturating photosynthetic photon flux density of 1400 µmol.m^-2^.s^-1^, air temperature of 23°C and reference VPD maintained between 1 and 1.4 kPa. Measurements began prior to the drying sequence to ensure the stabilization of gas exchange parameters inside the leaf chamber and to capture maximum *g*_s_. Parameters were logged every 60 sec and measurements lasted throughout the sequence until *g*_s_ reached constant basal values close to zero indicative of complete stomatal closure. As for leaf embolism, we then related each individual Ψ_leaf_ measurement (MPa) to the corresponding *g*_s_ measurement (see below for data analysis).

### 2.5. Growth and leaf nutritional status

All plants were regularly measured twice a week for basal stem diameter (± 0.01mm) before they were cut and used for the experiment; stem diameter was preferred over stem height as a more integrative indicator of growth to account for potential axillary growth. At the end of the drying sequence, the aerial biomass was partitioned between leaves from the main stem axis, leaves from sylleptic branches, woody branches and main stem. Plant material was dried in the oven at 105°C for 48 hrs and then weighed. The ratio between aerial woody biomass (main stem + branches) and total aerial biomass (woody + leaves) (R_woody_) was used as a proxy for aerial biomass partitioning between perennial and non-perennial compartments. The ratio between branch woody biomass and total woody biomass (R_branch_) was used as a proxy for branching. We also assessed leaf nutritional status by using a CCM-200 chlorophyll content meter (Opti-Sciences, Hudson NH, USA). Measurements were performed on three leaves, *i.e.* the one used for vulnerability measurements and the two leaves above and below. CCM readings were converted into Chlorophyll content (Chl) using the generic equation (excluding oak) reported in Brown et al. (2022).

### 2.6. Model fitting, parameter estimations and statistical analyses

Model fitting and statistical tests were all performed using the R software (R Core Team (2025). _R: A Language and Environment for Statistical Computing_. R Foundation for Statistical Computing, Vienna, Austria. https://www.R-project.org/).

Vulnerability curves were fitted using the following sigmoid function (Cochard et al. 2007): 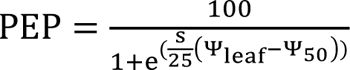 where PEP corresponds to the percentage of embolized pixels in the leaf lamina (%), Ψ_50_ corresponds to the water potential causing 50% of embolism (MPa) and *s* is the rate of embolism increase per unit drop in water potential (%.MPa^-1^). Stomatal response curves were analysed using the same function after transforming stomatal conductance into relative stomatal closure (%), enabling consistent parameter estimation across processes. Curves were all analysed using nonlinear mixed-effects models to account for repeated measurements within individuals and potentially unequal numbers of replicates. Models were fitted separately for each genotype and treatment combination using the *nlme* package, with individual identity included as a random effect. Besides Ψ_50_, the following physiological parameters were derived from fitted models: Ψ_12_ (the water potential inducing 12% of embolism, corresponding to the onset of cavitation), Ψ_88_ (the water potential inducing 88% of embolism, corresponding to hydraulic failure) and Ψ_gs90_ (the water potential inducing a 90% reduction in stomatal conductance, assumed to correspond to complete stomatal closure). Uncertainties around parameter estimates and fitted curves were quantified using parametric bootstrapping based on the variance–covariance matrix of the fitted models. Parameter sets were simulated (n = 1000) using functions from the *MASS* package and propagated to generate 95% confidence intervals for both predicted curves and derived parameters. Stomatal safety margins (SSMs) were defined as the difference between the point of stomatal closure (Ψ_gs90_) and vulnerability parameters (Ψ_12_, Ψ_50_, Ψ_88_, from less conservative to more conservative margins, respectively) and were calculated from bootstrap simulations to account for parameter uncertainty.

Differences between genotypes and treatments for parameter estimates (Ψ_gs90_, Ψ_12_, Ψ_50_, Ψ_88_, SSM_12_, SSM_50_ and SSM_88_) were interpreted based on confidence intervals. Differences between genotypes and treatments for measured traits such as π_o_, π_tlp_ and growth were analysed using one-way analysis of variance within genotypes or treatments. Tests were considered significant at *P*_value_ < 0.05.

Differences between genotypes in phenotypic plasticity were quantified using the relative distance plasticity index (RDPI). The RDPI was calculated for each trait as the absolute difference in phenotypic means between pairs of environments (C vs. N and C vs. K) divided by the sum of the phenotypic mean in the two environments and averaged across pairs: 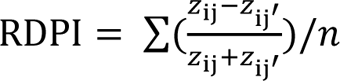 where z is the phenotype, i the genotype, j the environment, z_ij_ – z_ij’_ is the absolute distance for genotype i between environments j and j’, and n is the number of environment combinations (Arenas et al. 2025).

## Results

### Growth

Under control conditions, both genotypes showed comparable growth rate, chlorophyll content and biomass partitioning (*P* > 0.157) although PG-31 tended to be intrinsically more prone to branching than DRA-038 (Table 1). Additional N and K enhanced growth significantly in DRA-038 only (Table 1). Effects on biomass partitioning and chlorophyll content were not significant although additional N tended to increase branching and chlorophyll content, especially in DRA-038 (Table 1). Overall, treatment effects were genotype-dependant, DRA-038 benefiting most of the additional N and K supply.

**Table 1.** Growth, above-ground partitioning and leaf chlorophyll concentrations for the two genotypes of *Populus nigra* (DRA-038 vs. PG-31) in control (C), additional nitrogen (N) or additional potassium (K) treatments. Mean values (± SE).

| Genotype | Treatment | Diameter increment<br>(mm.day<sup>-1</sup>) | R<sub>woody</sub><br>(relative units) | R<sub>branch</sub><br>(relative units) | Chl<br>(g.m<sup>-2</sup>) |
| --- | --- | --- | --- | --- | --- |
| DRA-038 | C | 0.058 (0.003) | 0.48 (0.02) | 0.43 (0.10) | 0.52 (0.03) |
|  | N | 0.082 (0.012) | 0.47 (0.02) | 0.57 (0.14) | 0.58 (0.01) |
|  | K | 0.081 (0.006) | 0.51 (0.02) | 0.35 (0.02) | 0.59 (0.04) |
| PG-31 | C | 0.057 (0.006) | 0.49 (0.02) | 0.53 (0.06) | 0.46 (0.03) |
|  | N | 0.043 (0.007) | 0.48 (0.01) | 0.59 (0.05) | 0.47 (0.03) |
|  | K | 0.052 (0.006) | 0.50 (0.01) | 0.36 (0.18) | 0.51 (0.04) |

Diameter increment was computed for the timeframe between the onset of differential treatments and plant sampling. R<sub>woody</sub>, ratio between woody biomass (main stem + branches) and total aerial biomass (woody + leaves); R<sub>branch</sub>, ratio between branch woody biomass and the total woody biomass (main stem + branches); Chl, chlorophyll content estimated from CCM measurements (see Materials and Methods section).

### Leaf p-v parameters

Under control conditions, tests indicated no significant difference between genotypes in π_o_, π_tlp_, *RWC*_tlp_ and ε (*P* > 0.080) although PG-31 consistently exhibited lower π_o_, π_tlp_ and *RWC*_tlp_ values all indicative of an intrinsically higher tolerance to water deficit (Table 2). As for growth and leaf nutritional status, treatment effects were genotype-dependent and DRA-038 was the most responsive, although tests revealed no significant differences (*P* > 0.197; Table 2). Across all genotypes and treatments, π_o_ and π_tlp_ covaried positively (Fig. 1a).

**Figure 1.**
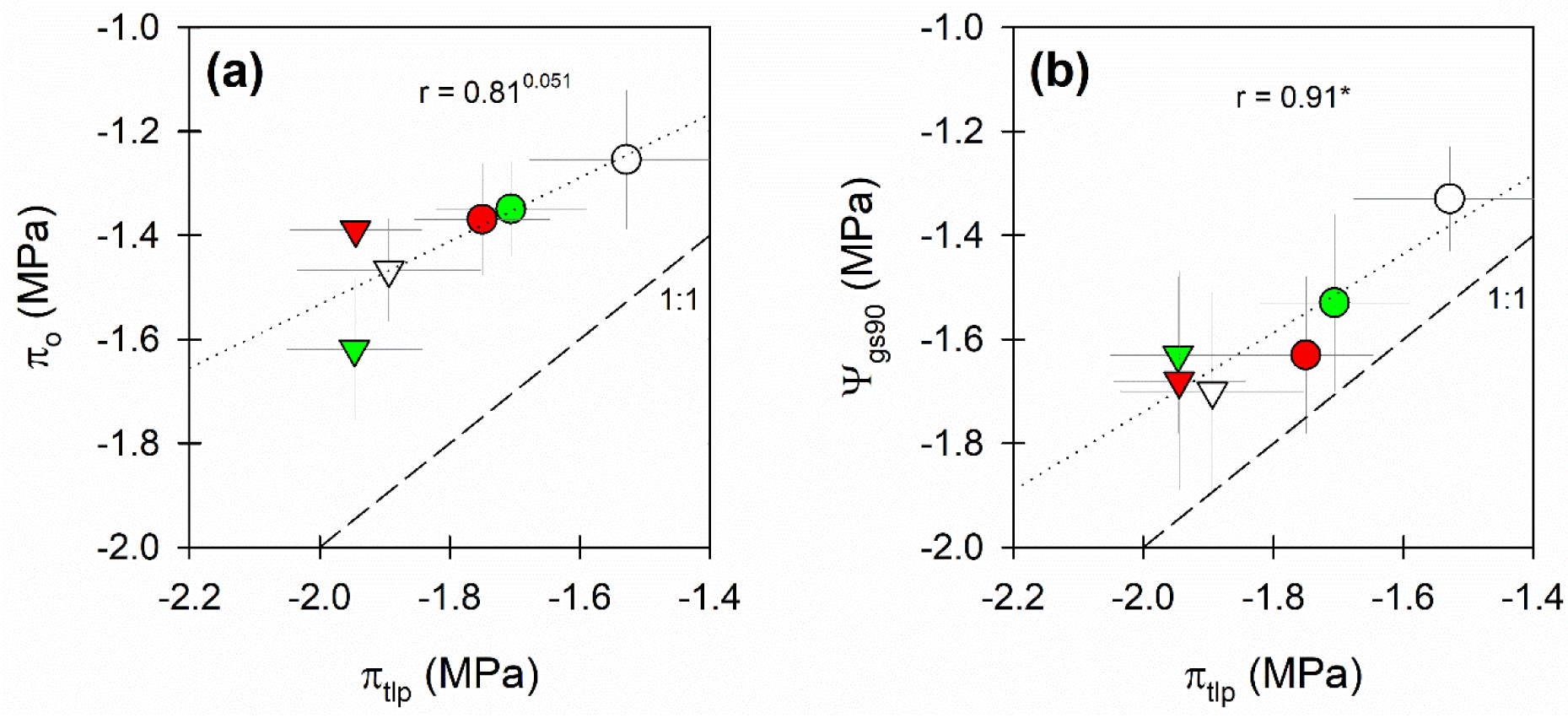
Coordination between bulk leaf osmotic potential at full hydration (π_o_), bulk leaf turgor loss point (π_tlp_) and the point of stomatal closure (Ψ_gs90_) across the two genotypes of *Populus nigra* and across control, additional N and additional K treatments (means ± SE for π_o_ and π_tlp_ and ± CI_0.95_ for Ψ_gs90_). Circles, genotype DRA-038; triangles, genotype PG-31; white symbols, control treatment; green symbols, additional N; red symbols, additional K. The dashed line corresponds to the 1:1 line; the dotted line corresponds to the linear regression across genotype and treatment means (r, Pearson’s correlation coefficient). Asterisks indicate significant correlations (*, 0.01 < *P* ≤ 0.05); superscripts indicate tendencies (0.05 < *P* ≤ 0.1).

**Table 2.** Pressure-volume (p-v) parameters derived from p-v curves on leaves of the two *Populus nigra* genotypes (DRA-038 vs. PG-31) in control (C), additional nitrogen (N) or additional potassium (K) conditions. Values are means (± SE) (*n* = 5-6).

| Genotype | Treatment | $\pi_o$ | $\pi_{tlp}$ | $RWC_{tlp}$ | $\varepsilon$ |
| --- | --- | --- | --- | --- | --- |
| DRA-038 | C | -1.25 (0.13) | -1.53 (0.12) | 87.21 (2.58) | 10.63 (1.44) |
|  | N | -1.35 (0.09) | -1.71 (0.12) | 85.53 (2.30) | 9.62 (2.25) |
|  | K | -1.37 (0.11) | -1.75 (0.10) | 87.90 (0.95) | 11.95 (1.80) |
| PG-31 | C | -1.47 (0.10) | -1.89 (0.14) | 83.92 (1.77) | 9.44 (1.47) |
|  | N | -1.62 (0.14) | -1.95 (0.12) | 85.77 (1.57) | 12.01 (2.04) |
|  | K | -1.39 (0.12) | -1.94 (0.10) | 82.43 (4.59) | 9.19 (1.24) |

$\pi_o$ , osmotic potential at full hydration (MPa); $\pi_{tlp}$ , turgor loss point (MPa), $RWC_{tlp}$ , relative water content at turgor loss point (%), $\varepsilon$ , modulus of elasticity (MPa)

### Stomatal conductance and stomatal closure

Mean maximal stomatal conductance (g_s_) under control conditions was not significantly different between genotypes (0.325 ± 0.033 vs. 0.303 ± 0.041 mmol.m^-2^.s^-1^ for DRA-038 and PG-31, respectively, *P* = 0.66). In both genotypes, N and K addition had no significant effect on maximal g_s_ (*P* > 0.067).

Following stem cutting, g_s_ declined rapidly (less than 10 min on average across all genotypes and treatments) as Ψ_leaf_ started to drop progressively (Fig. 2). Under control conditions, stomatal closure (Ψ_gs90_) occurred earlier in DRA-038 than in PG-31 (Table 3a). Both N and K addition tended to delay stomatal closure in DRA-038 while PG-31 was not affected by treatments (Table 3a). Stomatal closure systematically occurred before reaching the bulk leaf turgor loss point and values of Ψ_gs90_ covaried positively with values of π_tlp_ across genotypes and treatments (Fig. 1b).

**Figure 2.**
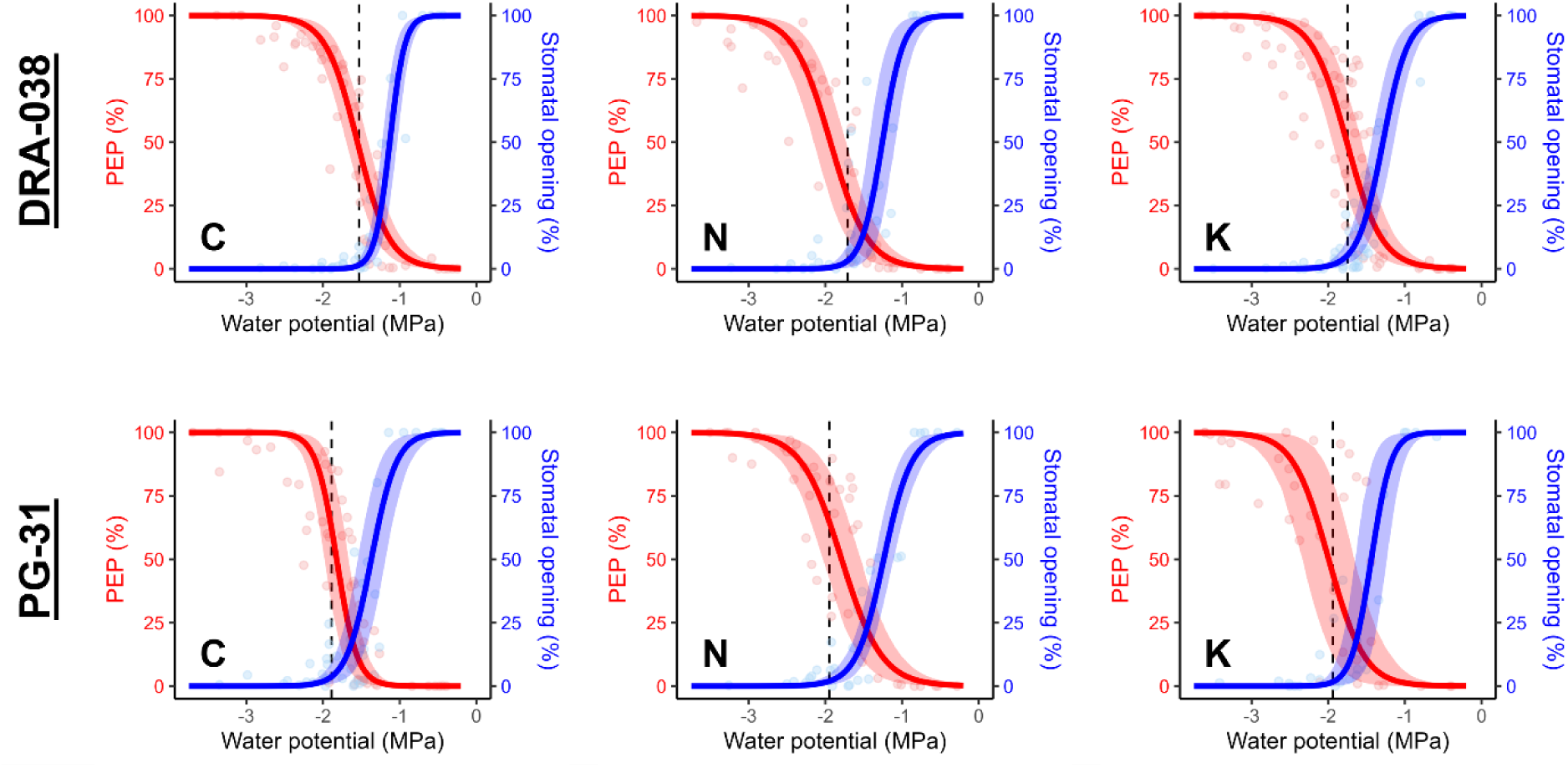
Stomatal and leaf embolism dynamics during drying in the two genotypes of *Populus nigra* (DRA-038 vs. PG-31) under control (C), additional N (N) or additional K (K) treatments. Leaf embolism was assessed using the optical vulnerability method and is computed as the % of embolized pixels (PEP). The dashed lines indicate the turgor loss point. Symbols correspond to individual data, and shaded areas correspond to 95% confidence intervals around response curves (see text for additional information).

**Table 3a.**
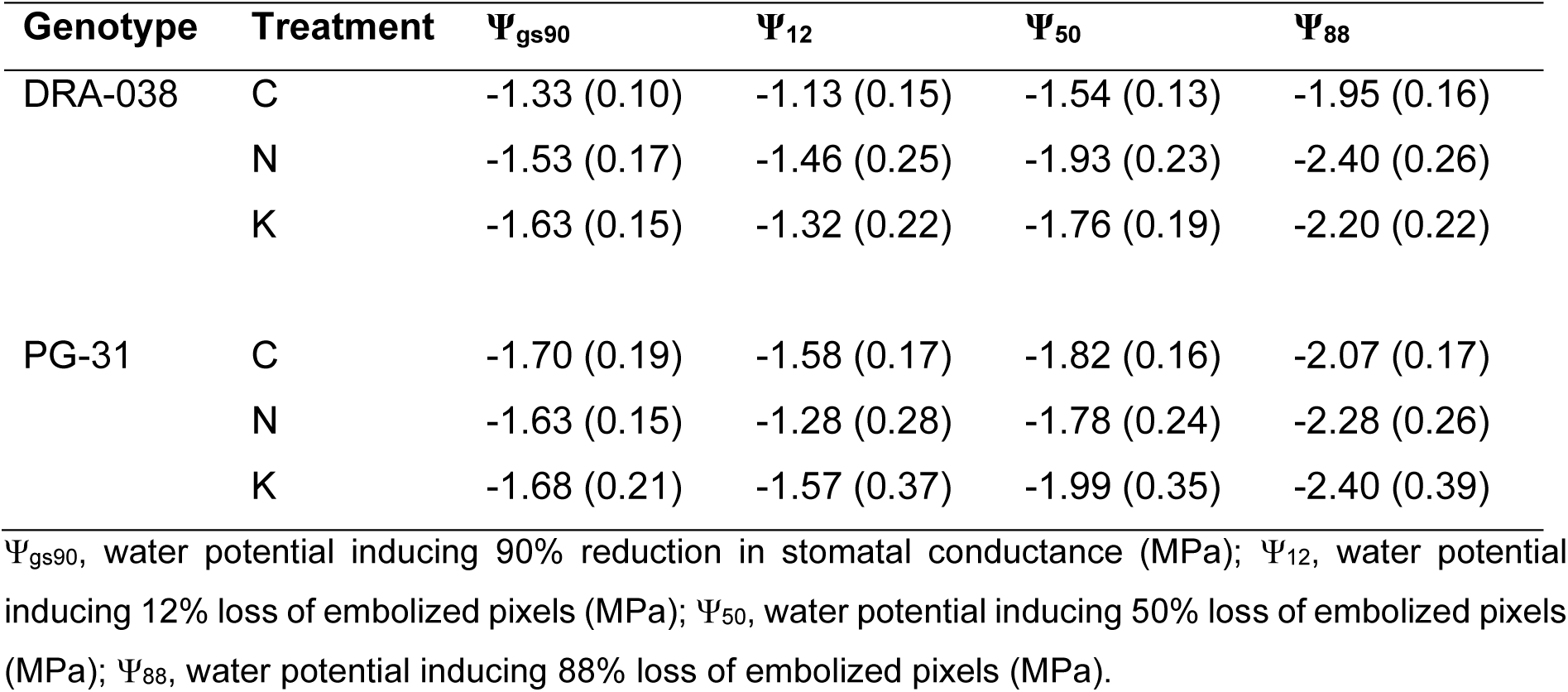
Stomatal closure point and vulnerability estimates for leaves of the two *Populus nigra* genotypes (DRA-038 vs. PG-31) in control (C), additional nitrogen (N) or additional potassium (K) conditions. Values are means (± SE) (*n* = 5-6).

### Leaf vulnerability to embolism and stomatal safety margins

The first events of xylem cavitation were observed following stem cutting most of the times within the first 30-60 min, with a maximum of 90 min, in all genotypes and treatments. The first main events typically occurred in the midrib and major veins before spreading across lower order veins (Fig. 3).

**Figure 3.**
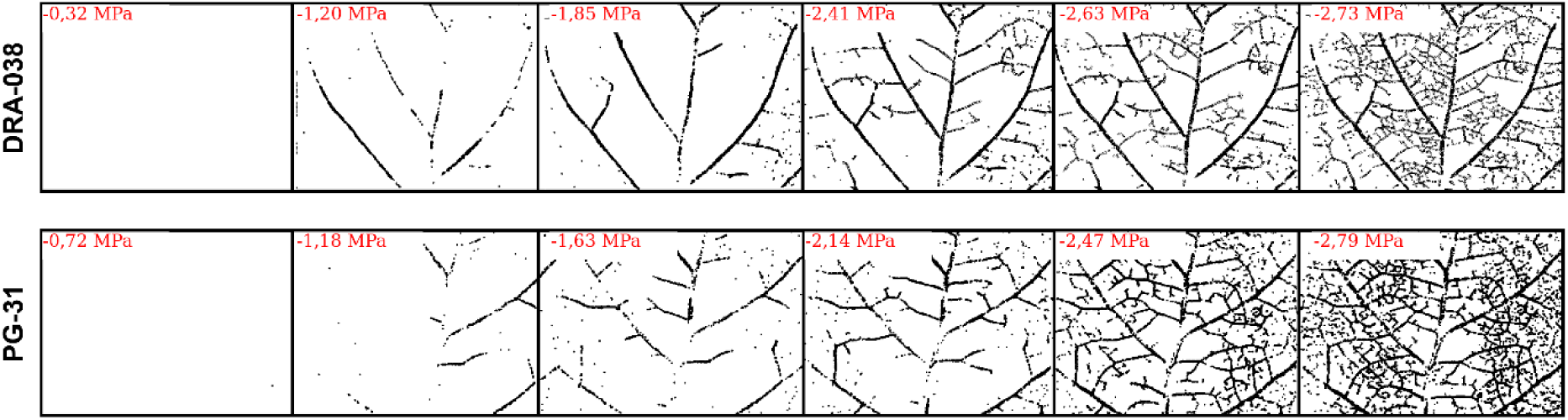
Examples of spatial and temporal dynamics of xylem embolism in leaves of the two *Populus nigra* genotypes studied (DRA-038 vs. PG-31) during drying. Images are from control treatment.

Vulnerability curves established from the percentage embolized leaf vein area showed sigmoidal shapes in all genotypes and treatments (Fig. 2). Values of Ψ_12_, Ψ_50_ and Ψ_88_ were highly correlated with no apparent offset (Sup. Fig. S2) indicating that genotypes and treatments primarily differed in intrinsic tolerance to embolism rather than in the slopes of vulnerability curves (Fig. 2). Under control conditions, leaves of DRA-038 were more vulnerable than leaves of PG-31 [Ψ_50-leaf_ = −1.54 MPa (CI_0.95_ = −1.41/-1.67 MPa) vs. −1.82 MPa (CI_0.95_ = −1.66/-1.98 MPa), respectively] (Table 3a). These differences matched those recorded on the main stem of the same genotypes in a previous independent experiment using the Cavitron technique [Ψ_50-stem_ = −1.62 MPa (CI_0.95_ = −1.52/-1.75 MPa) vs. −1.95 MPa (CI_0.95_ = −1.86/-2.04 MPa), respectively], stem values thus tending to be slightly more negative than leaf values. DRA-038 tended to be more responsive to N and K addition than PG-31 with systematically more negative vulnerability estimates than under control condition (Table 3a).

Mean values of SSM_12_ were slightly negative in all genotypes and treatments but CI_0.95_ overlapped with zero in most cases (Table 3b). Mean values of SSM_50_ were positive but also remained very low (0.12 to 0.40 MPa) and virtually not different from zero (Table 3b). Stomatal closure covaried positively with vulnerability estimates but not along the 1:1 line suggesting that SSMs may tend to increase with decreasing vulnerability to embolism (Fig. 4).

**Figure 4.**
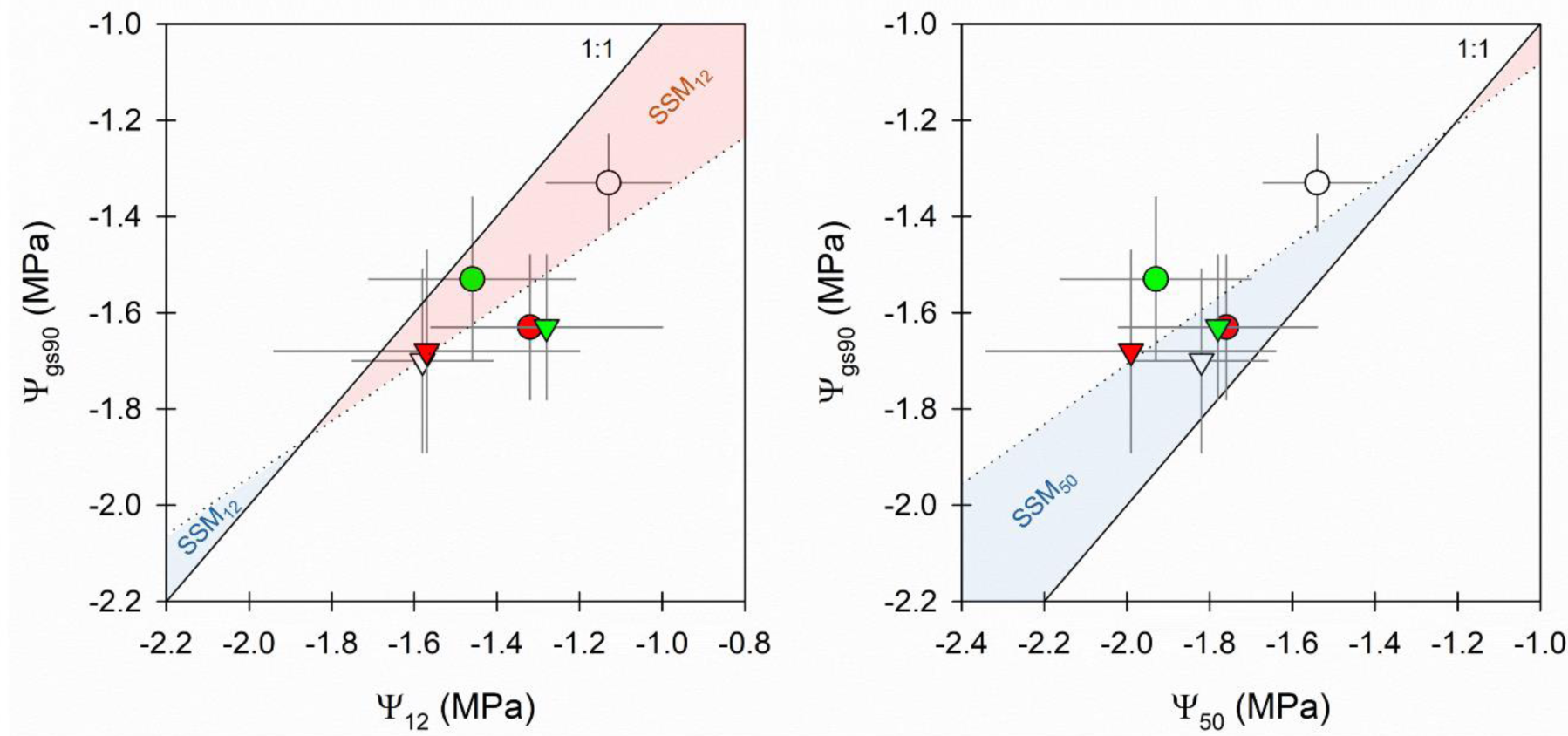
Coordination between the threshold for stomatal closure (Ψ_gs90_) and vulnerability to embolism estimates (Ψ_12_, Ψ_50_) (means ± CI_0.95_). Circles, genotype DRA-038; triangles, genotype PG-31; white symbols, control treatment; grey symbols, additional N; black symbols, additional K). The dashed line corresponds to the 1:1 line; the dotted line corresponds to the linear regression across genotype and treatment means. Projected safety margins (SSM_12_, SMM_50_) are shown in red when negative (i.e. Ψ_gs90_ more negative than Ψ_12_ or Ψ_50_) or in blue when positive (Ψ_gs90_ less negative than Ψ_12_ or Ψ_50_).

**Table 3b.** Stomatal safety margins (SSM, MPa) for the two genotypes of *Populus nigra* (DRA-038 vs. PG-31) in control (C), additional nitrogen (N) or additional potassium (K) treatments. SSMs were computed as the difference between Ψ_gs90_ and either Ψ_12_, Ψ_50_ or Ψ_88_ (SSM_12_, SSM_50_ and SSM_88_, respectively). Means (± CI_95%_).

| Genotype | Treatment | SSM<sub>12</sub> | SSM<sub>50</sub> | SSM<sub>88</sub> |
| --- | --- | --- | --- | --- |
| DRA-038 | C | -0.20 (0.18) | 0.21 (0.16) | 0.62 (0.19) |
|  | N | -0.07 (0.32) | 0.40 (0.29) | 0.87 (0.31) |
|  | K | -0.31 (0.27) | 0.13 (0.23) | 0.58 (0.25) |
| PG-31 | C | -0.12 (0.25) | 0.12 (0.25) | 0.37 (0.26) |
|  | N | -0.35 (0.37) | 0.15 (0.16) | 0.66 (0.30) |
|  | K | -0.11 (0.42) | 0.31 (0.32) | 0.72 (0.44) |

## Discussion

### The sequence of leaf traits and thresholds underlying leaf responses to water deficit in poplar

Severing shoots at the base allowed simulating progressive drought conditions over a relatively short period of time (Höchberg et al. 2017). Cavitation events were observed soon after stem cutting, typically within 30-90 min. Low order veins, i.e. midrib and secondary venation, were found to embolize first before cavitation progressively spread across higher order veins. This pattern was in line with previous reports in Angiosperm species displaying hierarchical, reticulate venation networks (Brodribb et al. 2016a,b; Höchberg et al. 2017; Avila et al. 2021). Reports of leaf vulnerability to embolism in poplar using the optical method remain limited (Mantova et al. 2023; Gonzalez et al. 2025) and sometimes not completely successful (Petruzzellis et al. 2023). Our findings indicated that the OV method can accurately resolve embolism dynamics in highly susceptible species such as poplars.

The values of embolism vulnerability recorded in our study (Ψ_50_ = −1.80 MPa across genotypes and treatments) aligned well with other reports on poplars (Fichot et al. 2015) including leaves measured with the OV method (Mantova et al. 2023; Petruzzellis et al. 2023; Gonzalez et al. 2025). The values obtained under control conditions indicated that the French genotype DRA-038 was more vulnerable than the Italian genotype PG-31 (Ψ_50_ = −1.54 MPa vs. −1.82 MPa, respectively). Most importantly, these values matched those obtained independently on stems of the same genotypes using the Cavitron reference technique with approximately the same range of variation (Ψ_50_ = −1.62 MPa vs. −1.95 MPa, respectively), although absolute values were slightly less negative with the OV method possibly related to organ differences (see discussion below). Our findings thus further indicated that the OV method can reliably resolve small differences in embolism vulnerability between closely related genotypes and can be deployed in the context of intraspecific studies (Dayer et al. 2020, 2022; Schell et al. 2026).

The coordination between leaf stomatal closure and embolism development has long been a matter of debate (Cochard & Delzon 2013). The development of reliable optical methods to directly visualize cavitation (Brodribb et al. 2016a) has, however, enabled unravelling the sequence with unprecedented high temporal resolution. Our findings fit with the consensus that significant cavitation does not occur before stomata close and that xylem embolism thus is not the trigger for stomatal closure (Brodribb et al. 2016a; Höchberg et al. 2017; Creek et al. 2020; Trueba et al. 2019; Blackman et al. 2019; Petek-Petrik et al. 2023). However, our findings also indicate that stomatal safety margins were virtually null in the studied poplars as SSM_12_ did not differ from zero in most cases while more conservative SSM_50_ remained very low too (0.22 MPa on average). To our knowledge, direct estimates of leaf SSMs with the OV method other than those reported in our study (i.e. directly measuring Ψ_gs90_, not using π_tlp_ as a proxy for Ψ_gs90_) are not available in poplars. However, other estimates of hydraulic safety margins in poplars (i.e. using proxies for stress intensity such as daily or minimum seasonal water potential instead of stomatal closure) typically converge towards small or even negative values, be it in leaves or in stems (Hukin et al. 2005; Fichot et al. 2010; Pan et al. 2016; Yu et al. 2025). Our findings show that poplars, at least *P. nigra*, do not operate at high levels of xylem embolism as sometimes assumed in such highly vulnerable species (Hukin et al. 2005; Secchi & Zwieniecki 2010) but do exhibit virtually null leaf stomatal safety margins.

Stomatal closure, and therefore the onset of embolism, systematically preceded bulk leaf turgor loss (by 0.21 MPa on average across genotypes and treatments) suggesting that guard cells response was to a certain extent uncoupled from bulk leaf water status. Although often assumed, the functional coordination between Ψ_gs90_ (or Ψ_gs50_) and π_tlp_ is labile across species (e.g. Brodribb & Holbrook 2003; Brodribb et al. 2003; Guyot et al. 2012). In Angiosperms, stomata can typically close in response to direct changes in water potential (passive hydraulic signal) as well as in response to chemical signalling such as abscisic acid (ABA) (ABA-mediated signal) (Buckley 2019). A role for an ABA-mediated signal was however very unlikely in our system because (i) dehydration was initiated by stem sectioning (no root-derived signal) and (ii) the very fast decline in stomatal conductance upon stem cutting (typically within 10 min max) was probably incompatible with *de novo* leaf ABA biosynthesis and signalling (McAdam et al. 2016; Brunetti et al. 2019). Instead, stomata most likely responded to a more localized hydraulic signal near the evaporation sites rather than to bulk Ψ_leaf_ (Guyot et al. 2012), possibly involving an early drop in the leaf hydraulic conductance outside the xylem pathways before significant cavitation even started (Scoffoni et al. 2017, 2023; Trifilò et al. 2021). An early stomatal closure is supposed to be a conservative mechanism, especially for highly sensitive species such as poplars, and agrees with the isohydric behaviour observed for the two genotypes studied here under prolonged moderate drought (Duplan et al. 2026).

Further along the dehydration sequence, π_tlp_ coincided more closely with Ψ_50_ (−1.79 vs. −1.80 MPa across genotypes and treatments, respectively) suggesting that bulk leaf loss of turgor in our poplar genotypes was reached only once significant embolism had already occurred. This was in line with a previous report on *P. nigra* (Trifilò et al. 2021) but contrasted with findings on *P. tremula × P. alba* where only 20% of embolism was predicted at π_tlp_ (Mantova et al. 2023), possibly reflecting differences among poplar species in their hydraulic design. Recent findings on grapevine also showed a good coincidence between π_tlp_ and Ψ_50_ (Dayer et al. 2020, 2022) while other studies on different species have reported a more variable coordination (Brodribb et al. 2003; Villagra et al. 2013; Trueba et al. 2019; Dayer et al. 2020, 2022), suggesting that drought tolerance at the cell level may be partly uncoupled from tolerance to embolism translating into possibly different drought response strategies, including within the genus *Populus*.

Early leaf shedding during water deficit has long been reported in poplar species and interpreted as an adaptive strategy (e.g. Braatne et al. 1992; Lu et al. 2010; Larchevêque et al. 2011; Bouyer et al. 2023). As such, leaves are considered as ‘fuses’ that may help prevent hydraulic failure of more costly perennial organs by mitigating further drops in water potential during the early stages of water deficit (Rood et al. 2000; Lu et al. 2010; Wolfe et al. 2016). Interestingly, our findings indicated a slightly higher vulnerability to embolism in leaves compared to stems, suggesting a possible vulnerability segmentation that might promote early leaf shedding (Höchberg et al. 2017), although we acknowledge our study was not initially designed to test this directly (i.e. independent experiments, different techniques). Vulnerability segmentation, where distal organs such as leaves and petioles are more vulnerable than stems and/or roots, is not a general rule (e.g. Charrier et al. 2016; Levionnois et al. 2020; Avila et al. 2021; Li et al. 2020b), including in poplar (e.g. Huber et al. 2023; Wilkening et al. 2023; Rimer et al. 2025). Nevertheless, even in the absence of segmentation, the most peripheral parts of the plants are still expected to endure the most negative water potentials and to cavitate first (Wolfe et al. 2016). Model simulations have indicated that reducing leaf area is the most effective physiological adjustment to delay the time to hydraulic failure in poplar (Lemaire et al. 2021). Although leaf loss is costly, pioneering species such as poplars remain on the acquisitive side of the species spectrum and are typically able to resprout rapidly from roots or stem axillary buds as long as meristem integrity is preserved (Lu et al. 2010). The combination of near-zero stomatal safety margins with high leaf minimum conductance as already reported in poplars (Grünhofer et al. 2022; Garen & Michaletz 2025), sustaining residual water loss despite stomatal closure, should result in a very short timeframe between stomatal closure, massive xylem hydraulic failure and ultimately cell death. Such a hydraulic design, besides possible vulnerability segmentation, likely results in an early hydraulic disconnection of leaves in the case of severe water stress, acting as an efficient means to delay hydraulic failure in perennial compartments. The typical progression of leaf yellowing and senescence in poplar under progressive water deficit, i.e. from the base to the top, suggests that basal leaves may be more vulnerable to embolism than apical leaves as observed in grapevine (Höchberg et al. 2017), but this deserves to be formally tested.

### Plasticity and coordination of leaf traits and thresholds across genotypes and treatments

While the two *P. nigra* genotypes displayed comparable growth rates under control conditions despite contrasting geographic origins, differences were more readily apparent when considering the plastic responses to N and K treatments. Additional N and K both stimulated secondary growth in the genotype DRA-038 only. However, despite a similar outcome, the underlying effects of N or K treatments on above-ground resource allocation in DRA-038 were different. Additional N indeed tended to increase branching, although without affecting allocation to leaves relative to woody parts, while additional K did not. Sylleptic branch development is known to respond rapidly to increased N availability as a result of coordinated shifts in C and N metabolisms (Cooke et al. 2005; Novaes et al. 2009) and is a marker of increased biomass production in young poplar saplings (Rae et al. 2004). Alternatively, although K availability can also have a major influence on growth and allocation in fast growing tree species such as eucalypts (Epron et al. 2011, 2015; Mateus et al. 2022), the effects remain comparatively less documented in poplar (Ache et al. 2010). Overall, our findings indicated that N and K addition stimulated growth (1) in a genotype-dependent manner, possibly reflecting different population life histories regarding local nutrient availability, and (2) through distinct underlying modifications of sink-source relationships.

Although plasticity in leaf hydraulic traits was typically more restrained than growth (probably explaining why significant treatment effects were also more difficult to record), the patterns between genotypes remained largely comparable, i.e. the genotype DRA-038 was the most responsive to treatments (Fig. 5). Additional N and K decreased Ψ_gs90_, π_tlp_ and Ψ_50_ in DRA-038 while PG-31 remained globally unaffected. In other words, the intrinsically less tolerant genotype under control conditions (DRA-038) turned out to be also the most plastic in response to increased N and K availability. Recent findings on the same two poplar genotypes have revealed that the genotype DRA-038 also displayed exacerbated hormonal and transcriptional responses in cambium-derived tissues when subjected to moderate water deficit while PG-31 exhibited much more stable responses (Duplan et al. 2026), confirming differences between genotypes in terms of plasticity but at a different scale and in response to a different cue. Other studies have reported significant associations between basal trait values and plastic responses for Ψ_50_ specifically in response to moderate water deficit in different poplar genetic backgrounds (see Fichot et al. 2015), suggesting the occurrence of a global trade-off between intrinsic tolerance to water deficit and plastic capacities. The fact that the nutrient-induced leaf hydraulic adjustments observed in the genotype DRA-038 in our study were mirrored by changes in growth suggests that this trade-off may be at least partially mediated by differences in growth strategies underpinned by local adaptation to resource availabilities (Fichot et al. 2024).

**Figure 5.**
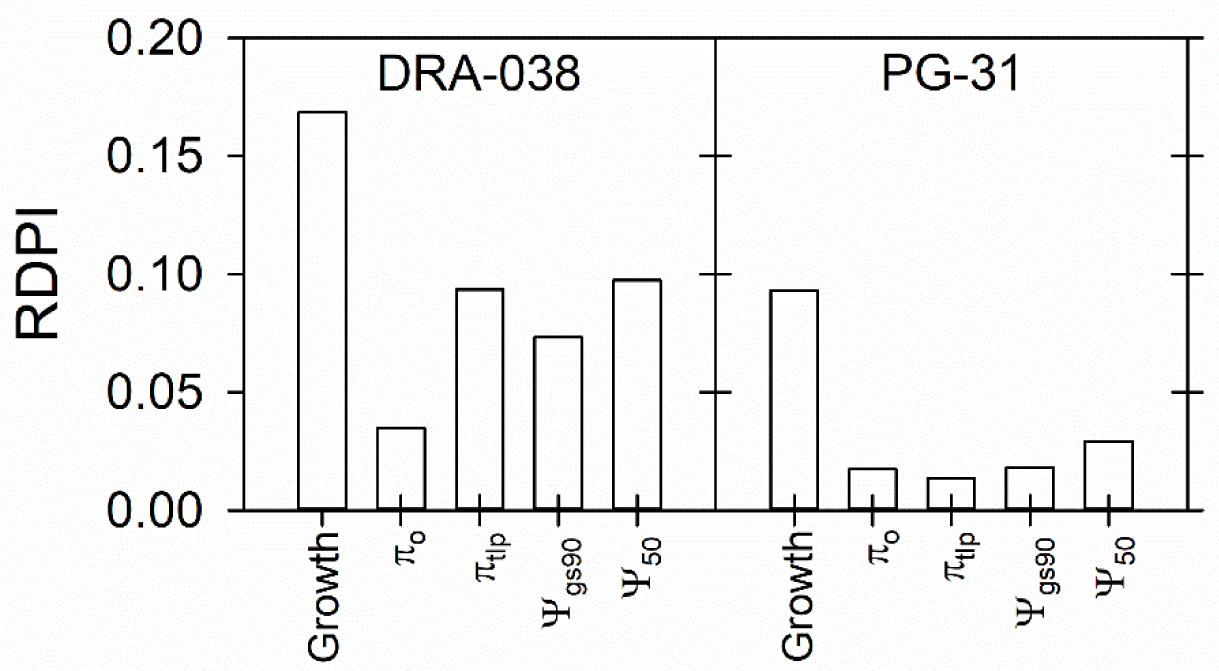
Comparison of phenotypic plasticity between the two *Populus nigra* genotypes (DRA-038 *vs*. PG-31) averaged across environments for growth and leaf traits involved in the sequence of response to water deficit using the relative distance plasticity index (RDPI). Growth corresponds to diameter growth rates. π_o_, bulk leaf osmotic potential at full hydration; π_tlp_, bulk leaf turgor loss point, Ψ_gs90_, point of stomatal closure, Ψ_50_, leaf water potential inducing 50% xylem embolism.

Adjustments in leaf p-v parameters and stomatal functioning in response to nutrient availability have already been reported, although effects and magnitude can be largely species- or even genotype-dependent (e.g. Domec et al. 2009; Villagra et al. 2013; Villar-Salvador et al. 2013; Battie-Laclau et al. 2014; Fang et al. 2018; Beikircher et al. 2019; Zhang et al. 2021). In our study, increased N and K availability both delayed stomatal closure by 0.2-0.3 MPa in the plastic genotype DRA-038, without affecting maximum leaf gas exchange rates. Interestingly, additional N and K also decreased π_tlp_ with the same magnitude and, to a lesser extent, tended to decrease π_o_. This suggested that when stomatal dynamics was responsive to treatments, this mainly occurred through changes in cell turgor-related status that were primarily driven by osmotic adjustments (Bartlett et al. 2012).

Vulnerability to drought-induced embolism can also be responsive to changes in environmental conditions provided the time-lapse for acclimation is sufficient to produce newly acclimated xylem conduits (Fichot et al. 2015). Owing to the continuous growth habit of poplars, we ensured that the leaves sampled for vulnerability measurements in our study were systematically mature and had been produced during the period of nutrient treatments such that the propension to acclimation was maximized. Once again, additional N and K both decreased leaf vulnerability to embolism (Ψ_50_) in the genotype DRA-038 by 0.39 and 0.22 MPa. The marginal effect of additional K was in line with other reports (Harvey & van den Driessche 1999; Mateus et al. 2024). The effect of additional N was more surprising considering that increased N availability has been frequently associated with slight increases in vulnerability to embolism in poplar stems (Hacke et al. 2010; Plavcová & Hacke 2012; Bouyer et al. 2023). However, findings in other species have been mixed (Bucci et al. 2006; Villagra et al. 2013; Fang et al. 2018; Beikircher et al. 2019; Zhang et al. 2021; Fan et al. 2022) and may depend on organs.

The most direct outcome of our findings was that, despite inherent genetic variation in trait values and/or in trait plasticity, the sequence of leaf traits and thresholds involved in the response to water deficit remained tightly coordinated across genotypes and treatments. In our model system, an increased leaf capacity to tolerate wilting thus translated into delayed stomatal closure and delayed cavitation occurrence. As a consequence, SSMs remained largely stable, in line with the homoeostasis already reported in the hydraulic safety margins of Mediterranean trees (Moreno et al. 2024) and constrained to near-zero values in the case of our poplars, although projections suggested that higher tolerance to embolism might be associated with slightly higher SSMs (Fig. 4) as observed across species (Martin-StPaul et al. 2017). Similar coordination between leaf-level hydraulic traits within species across genetic units has been reported in grapevine, although SSMs were found to be variable in this specific case (Dayer et al. 2020, 2022), but evidence of such coordination across varying environments remains so far undocumented to our knowledge. Our findings thus contribute to fill this gap, confirming an inherent genetic basis but also revealing a strong functional mechanistic basis over trait coordination. The coordination between the dynamics of stomatal closure and bulk leaf water relation parameters derived from p-v analyses somehow can be expected since stomata mechanistically respond to water status, although not necessarily bulk leaf water status *per se* (Buckley 2019). In contrast, the mechanistic rationale for a functional linkage with vulnerability to embolism (i.e. beyond simple trait co-variation or co-selection), which is inherently determined by the ultrastructural properties of xylem interconduit pits (Choat et al. 2008), remains to be elucidated.

## Conclusion

Despite inherent genotypic variation and/or phenotypic plasticity in traits, the sequence of leaf hydraulic thresholds involved in the response to water deficit remained tightly coordinated in the riparian species *Populus nigra*, supporting a strong mechanistic link between stomatal regulation, cell water status and xylem integrity across genetic units and varying environments. As a consequence, SSMs remained largely stable and constrained to near-zero values, questioning both the physiological relevance of such small variations in hydraulic safety margins and, from a practical standpoint, the feasibility of exploiting them in poplar breeding. We acknowledge that the rapid dehydration imposed by stem cutting in our system most likely precluded any contribution of ABA-mediated signaling to stomatal closure; thus, whether progressive, field-realistic drought would alter the relative timing of thresholds reported here deserves to be tested although findings on grapevine have already indicated a good agreement (Höchberg et al. 2017). Our findings further support the occurrence of a trade-off between basal trait values and the extent of phenotypic plasticity for hydraulic traits in poplars, calling into question which kind of ideotype should be favoured in the future, i.e. inherently more tolerant but less plastic vs. inherently less tolerant but more plastic. Additional work explicitly addressing how these trade-offs relate to performance and fitness in combination with model-assisted ideotyping framework (Dayer et al. 2022) should help answer these questions, especially in interspecific hybrids that can exhibit distinctive trait patterns (Fichot et al. 2010) and that form the core of poplar planting material.

## Acknowledgements

This work received funding from the Région Centre-Val de Loire (CPER ValoPat 2021-2027, project ECOCOPHYSI-O 2021-146895). The preliminary experiment (stem vulnerability to embolism) was performed in the framework of the research project EPITREE (ANR-17-CE32-0009-01) and we thank Prof. S. Maury (Université d’Orléans, INRAE USC1328, UR 1207 Physiology Ecology and Environment, P2e) for his support and scientific discussions on the physiological and molecular responses of the two studied genotypes; we also thank Dr. H. Cochard (UMR PIAF, INRAE-Université Clermont Auvergne) for preliminary discussions and access to Cavitron facilities (Phénobois, INRAE, 2018. Plateforme de Phénotypage de Propriétés Hydrauliques et Physico chimiques du Bois de Ressources Génétiques Ligneuses, https://doi.org/10.15454/1.5572410490640864E12) as well as Dr. V. Segura (UMR AGAP Institut, Univ Montpellier, CIRAD, INRAE, Institut Agro Montpellier) for his help in selecting the six original genotypes based on genetic and phenotypic data. We finally thank P. Poursat (INRAE Experimental Unit GBFOR, INRAE, 2018, Forest Genetics and Biomass Facility https://doi.org/10.15454/1.5572308287502317E12) and the PNRGF Guémené-Penfao for providing tree cuttings.

## Author contributions

DC set up the experiment, participated in all measurements and data analyses, and helped revising the manuscript; LB participated in preliminary tests and methodological development; ILJ produced plant material and helped with the experimental setup; RF designed the experiment, participated in measurements, analysed data and wrote the first draft of the manuscript; all authors approved the final version of the manuscript.

## Conflicts of interest

The authors have no conflict of interest to declare.

## Data availability

Data will be made available upon request to the authors

## Supplementary data

**Fig. S1.**
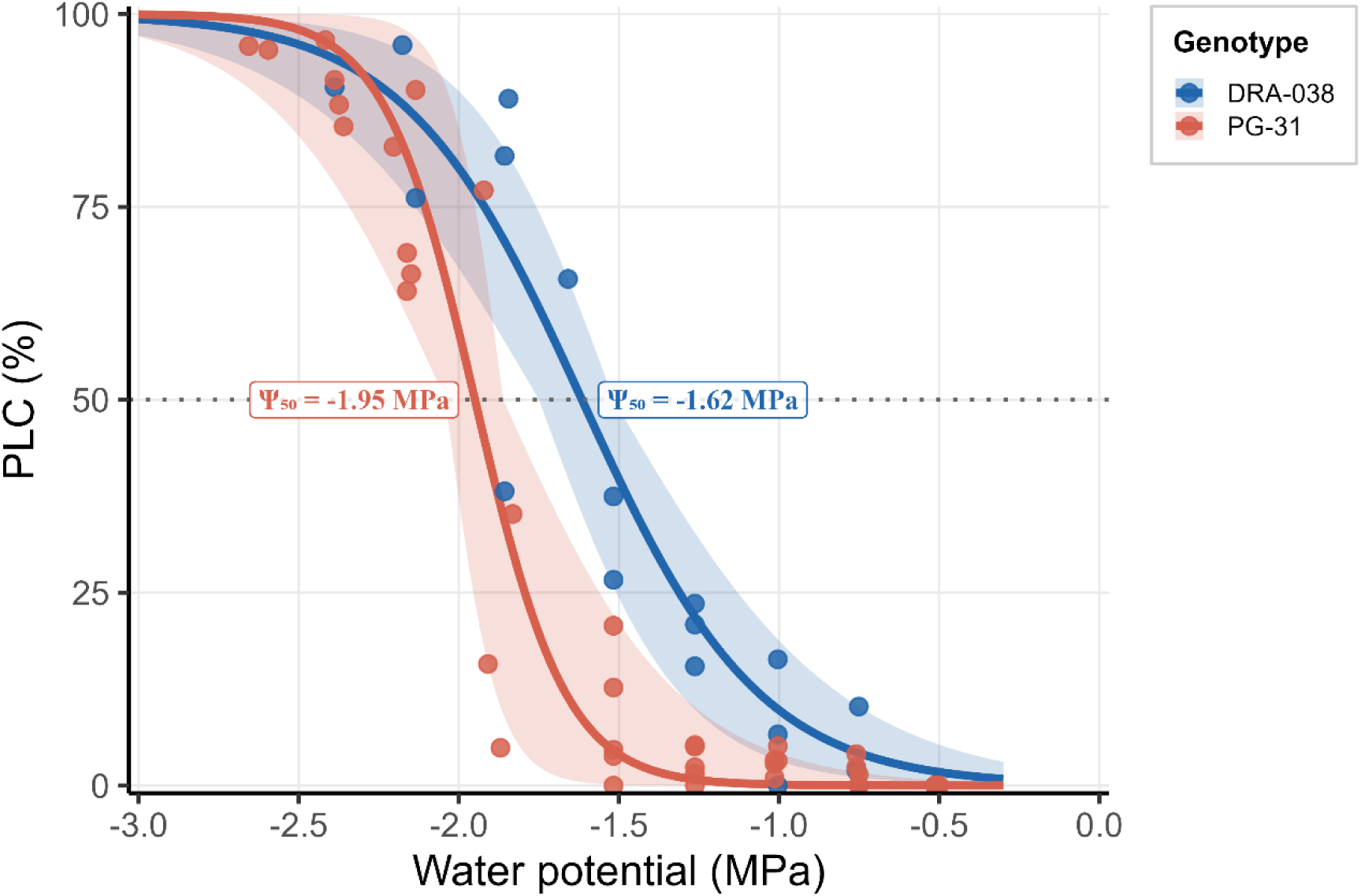
Stem vulnerability curves for the two *Populus nigra* genotypes (DRA-038 vs. PG-31) obtained with the Cavitron technique. Measurements are from an independent, preliminary experiment conducted on cuttings grown under control greenhouse conditions. A sigmoidal function was fitted to individual data of each genotype using the *fitplc* package and represented using the *ggplot2* package in R (R Core Team, 2025). Symbols correspond to individual data (n = 3-5 trees per genotype) and envelopes correspond to confidence intervals at 95% calculated from bootstrapping (Duursma & Choat, 2017). PLC, percent loss in hydraulic conductance; P_50_, water potential inducing 50% loss in hydraulic conductance.

**Fig. S2.**
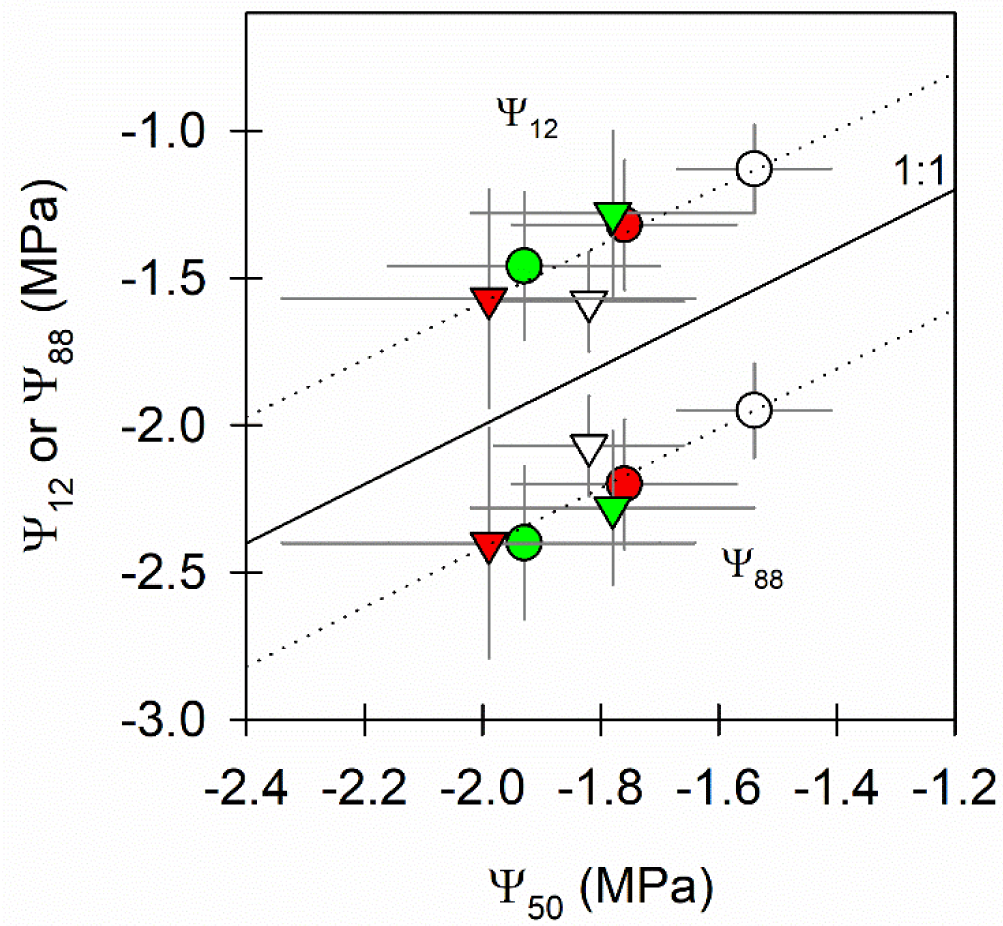
Relationships between vulnerability estimates (Ψ_12_, Ψ_50_, Ψ_88_) across the two *Populus nigra* genotypes (DRA-038 = circles, PG-31 = triangles) under control (white symbols), additional N (green symbols) and additional K (red symbols) conditions. Data represent means ± CI_0.95_ (n = 4-6 per genotype and treatment) and were obtained using the optical vulnerability (OV) method. Dashed lines represent linear regression between either Ψ_12_ or Ψ_88_ and Ψ_50_. The solid line indicates the 1:1 line.

## Notes

### Competing Interest Statement

The authors have declared no competing interest.

